# The Modular Adaptive Ribosome

**DOI:** 10.1101/068361

**Authors:** Anupama Yadav, Aparna Radhakrishnan, Anshuman Panda, Amartya Singh, Himanshu Sinha, Gyan Bhanot

**Author notes:** **Authors for correspondence**: Himanshu Sinha, Department of Biotechnology, Indian Institute of Technology Madras, Chennai 600036, India; and Gyan Bhanot, Department of Molecular Biology and Biochemistry, Rutgers University, Piscataway, New Jersey 08854, USA.

## Abstract

The ribosome is an ancient machine, performing the same function across organisms. Although functionally unitary, recent experiments suggest specialized roles for some ribosomal proteins. Our central thesis is that ribosomal proteins function in a modular fashion to decode genetic information in a context dependent manner. We show through large data analyses that although many ribosomal proteins are essential with consistent effect on growth in different conditions in yeast and similar expression across cell and tissue types in mice and humans, some ribosomal proteins are used in an environment specific manner. The latter set of variable ribosomal proteins further function in a coordinated manner forming modules, which are adapted to different environmental cues in different organisms. We show that these environment specific modules of ribosomal proteins in yeast have differential genetic interactions with other pathways and their 5’UTRs show differential signatures of selection in yeast strains, presumably to facilitate adaptation. Similarly, we show that in higher metazoans such as mice and humans, different modules of ribosomal proteins are expressed in different cell types and tissues. A clear example is nervous tissue that uses a ribosomal protein module distinct from the rest of the tissues in both mice and humans. Our results suggest a novel stratification of ribosomal proteins that could have played a role in adaptation, presumably to optimize translation for adaptation to diverse ecological niches and tissue microenvironments.

## INTRODUCTION

A single celled organism displays a range of phenotypes to survive in diverse environments. In complex multicellular organisms, in addition to the external environment, tissue specific cell types display specialized mechanisms to regulate phenotype in local tissue environments. Much of the research in biology has been directed towards understanding the basis of the information flow that gives rise to these diverse phenotypes. This has resulted in the identification of many regulatory processes [1,2] which fine-tune transcriptional expression and modulate the translation of mRNA into proteins [3,4] in response to external and environmental or tissue specific signaling cues. However, in spite of its essential role in this cellular information flow, the ribosome has always been regarded as an inert participant in the information flow that regulates cellular and tissue states.

One may ask, “Why is it that, in spite of tantalizing clues to the contrary, this belief in an invariant, environment independent ribosome has not been significantly challenged?” A possible reason might be that the high degree of conservation of ribosomal proteins across the three domains of life, viz., Archaea, Bacteria and Eukarya, and the slow evolution rates of ribosomal protein sequences [5] seem sufficient evidence for an essential, invariant ribosome, which plays no regulatory role in cells and tissues or in adaptation and speciation.

However, thermophiles have ribosomes that function at extreme temperatures, suggesting that the ribosome has been adapting to the environment for over 3 billion years. There are different types of ribosomes in each domain of life, and additionally, the mitochondrial ribosome distinct in composition from the cytosolic ribosome [6]. It is therefore worth investigating whether the cytosolic ribosome has also evolved ways to optimize its composition in response to environmental cues. Testing the ribosomal composition at a protein level is technically challenging, but significant evidence has been accumulating from the study of ribosomal proteins that argues for a variability in the composition of the ribosome [7-9]. Deletion experiments in yeast show that different ribosomal proteins have a differential effect on replicative lifespan [8]. Transcriptional studies in mice [7] and humans [9] show tissue specific expression of ribosomal proteins. Specific ribosomal proteins are known to be associated with different types of cancers [10,11] and mutations in specific ribosomal proteins result in a class of disorders called ribosomopathies [12,13]. Whereas all these mutations have common effects across development, some cause developmental disorders in specific tissues [12,13], suggesting tissue specificity of the function of at least some ribosomal proteins.

To date, most of these effects have been ascribed to extra-ribosomal functions of ribosomal proteins [14,15]. However, at least some specialized translation by the ribosome is controlled by specific ribosomal proteins [7,16,17], showing that the extra-ribosomal nature of such control is not generic. While preliminary, these studies suggest that ribosomes with variable components may exist to optimize translation, depending on environmental and signaling cues [18-20]. Analysis of the stoichiometry of ribosomal proteins in yeast and embryonic stem cells has demonstrated differential protein composition of ribosomes in different conditions [21], further substantiating the possibility of ribosomal variability at a protein level.

In this paper, we question the unitary nature of the ribosome within an organism across environments by analyzing different properties of ribosomal proteins. Does the observed environment and tissue specific variation in ribosomal proteins represent regulation of the ribosome that is important in evolution and or adaptation? In a changing environment, do all ribosomal proteins have similar properties, or are specific ribosomal proteins used in an environment dependent manner? In this paper we begin to address these questions by uncovering evidence from data analyses of yeast deletion [22] and interaction [23] datasets, and the ENCODE [24] and GTEx [25] expression datasets of mice and humans.

## MATERIALS AND METHODS

### Yeast Data

Growth data and associated microarray files for a genome-wide yeast homozygous deletion collection [22] for all environments were downloaded from http://chemogenomics.stanford.edu/supplements/global/download.html. The normalized gene intensity values from the microarray data were used for analysis. The data of the replicates for each environment were collated using their median. The intensity values were scaled to zero mean and unit variance across all genes and then compared between the rich growth (YPD) and a stress condition, as well as between the pairs of stress conditions (S1 Table). A set of 68 non-essential structural ribosomal proteins with microarray tag intensity greater than 2 standard deviations from the background intensity was used for the analysis.

Thirty four yeast-specific pathways, known to be involved in growth and stress resistance, described in Wikipathways database [26] were downloaded from http://www.wikipathways.org/index.php/WikiPathways (S2 Table).

### Clustering of genetic interactions in yeast

Phenotype data of pair-wise quantitative genetic interactions of 1,712 yeast genes derived from SGA [23] dataset was downloaded from http://drygin.ccbr.utoronto.ca/index.html along with their genetic interaction and *P* values. Genetic interactions of intermediate stringency (*P* < 0.05) were selected for our analysis.

In order to cluster the genes into different genetic clusters based on their double-deletion interactions, an adjacency matrix with the interaction scores for the genetic interactions was created from the above data. Pearson correlation among the genes was calculated based on their double deletion interaction scores. Genes with *r*^*2*^ > 0.2 were grouped in the same cluster (S2 Table). The Markov Cluster (MCL) algorithm [27] was applied, using R package function *mcl*, to get highly connected clusters of genes, with an inflation factor of 1.4, as used by the original study Costanzo et al.[23].

### Analysis of clusters and pathways in yeast

MCL clusters with 5 or more genes were considered for further analyses (S2 Table). The F-test statistic, BF-test statistic [28], *P* values and variances were computed on the standardized phenotype data derived from the above-standardized Hillenmeyer dataset. Genes in each cluster in each condition were compared to their respective YPD control condition. An identical analysis was carried out for each of the thirty-four pathways downloaded from Wikipathways [26]. Only pathways with 5 or more genes were used (S2 Table). A total of 90 MCL clusters and pathways were considered further and they were associated with several functional categories relevant for growth in yeast, such as known biochemical pathways (TCA, glycolysis), signaling pathways (PKA, MAPK), protein complexes (ribosome, proteasome) and a large number of various other genetic networks (S2 Table).

### Hierarchical clustering of cytoplasmic ribosomal proteins in yeast

The genes coding for cytoplasmic ribosomal proteins (n = 68) were considered in 26 environments of the above-standardized Hillenmeyer dataset. A hierarchical clustering of both the environments and the genes were carried out using a Euclidean distance metric using the data, which was standard normalized (mean zero, variance unity) across the ribosomal protein genes.

### Enrichment of genetic interactions in Clusters A, B and C

Enrichment of GO categories was carried out in the 121 genes identified to interact with at least 10 out of 65 ribosomal proteins (S3 Table). These 121 interactors interacted either positively or negatively with ribosomal proteins, with a few genes showing both positive and negative interactions (S3 Table). To identify genetic interactors specific to each Cluster, genes interacting with >10% of the ribosomal proteins in each cluster were identified.

### Phenotyping single deletions of yeast ribosomal proteins and their deletions with *GCN5* deletion in various environments

Single ribosomal protein and regulatory gene deletions of S288c background (BY4741) were obtained from the haploid genome-wide deletion collection library [29] (GE Healthcare Dharmacon Inc.). Additional double-gene deletions were generated using a previously described protocol [30]. SK1 wild type and deletion strains were obtained from Wilkening et al. [31]. Strains were phenotyped in YPD, YPD+Menadione (50µM) and YPD+CdCl_2_ (10µM). Spot dilutions, ranging from 10^−3^ to 10^−8^ dilutions were incubated at 30°C and phenotyped at 24, 36 and 48h. The strain and primer list is given in S4 Table.

### Sequence analysis of coding and 5’UTR regions in SGRP collection

*S. cerevisiae* and *S. paradoxus* ribosomal protein and control gene sequences from the SGRP strains [32] were downloaded from http://www.moseslab.csb.utoronto.ca/sgrp/blast_original. Sequence alignments, estimation of the maximum likelihood tree (1,000 permutations) and nucleotide diversity were performed using MEGA 6.06 [33] with default parameters (S5 Table).

For the three gene clusters based on the interaction modules, the nucleotide diversity in the coding and promoter regions of the genes in each cluster was evaluated using Shannon Entropy function as a metric. The sequences of the 5’UTR and coding regions of each gene corresponding to the different strains were aligned separately using the software MUSCLE [34] with default parameters. In the aligned set of sequences for each gene, the mutated sites were identified, and corresponding to each kind of base at the given site, were assigned a value,

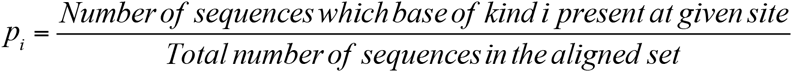

Here *i* = 1, 2, 3, 4 corresponds to A, T, G, C respectively.

The Shannon Entropy at each mutation site was computed using the standard definition:

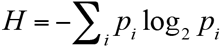

Variations within a given sequence set can occur in two principal ways: (i) variations in the proportion of bases at any given mutation site, and (ii) variations in the number of sites of mutations. The simplest quantity that accounts for both these is the sum of the *H* values for all the mutation sites. A higher value for the sum would indicate greater nucleotide diversity in the corresponding 5’UTR or coding region respectively for each gene across the different strains. To eliminate any length bias in the comparisons, the sum of the *H* values was normalized by the length of the aligned sequences for each 5’UTR or coding region (S5 Table).

### Prediction of transcription factor binding sites

The database YEASTRACT [35] was used to download known transcription factor (TF) binding sites of the various ribosomal proteins and to predict potential binding sites on UTR of ribosomal proteins of different SGRP strains. Of the 700 TFs reported in YEASTRACT, 216 have been shown to experimentally bind to the promoter region of at least one ribosomal protein present in the three Clusters identified in our study (S6 Table). These 216 TFs were enriched for various signaling pathways and chromatin remodeling complexes (S6 Table), substantiating the enrichment of chromatin remodelers among the positive genetic interactors of ribosomal proteins. Of these 216, the TFs binding exclusively to ribosomal proteins in Cluster A, B and C were identified.

### Analysis of human and mouse ENCODE and GTEx data

Tissue specific count (transcripts per million or TPM) data for human and mouse were downloaded from ENCODE (https://www.encodeproject.org/), and mapped to Entrez genes using annotation packages *org.Hs.eg.db* and *org.Mm.eg.db* in R. The genes that were not expressed in any replicate were discarded. Replicates in which an unusually high (as determined from *sfigx_h* and *sfigx_m*) fraction of Entrez genes were not expressed were discarded as well. In the remaining replicates, to reduce relative systematic error among replicates, the median for each gene was normalized to unity in each replicate by dividing the count for the gene by the median count in each replicate array. These median adjusted TPM values were log transformed to obtain the final expression *X* of each gene in each replicate as follows:

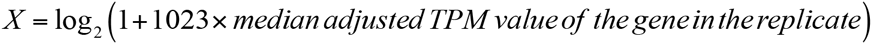

This normalization ensured that genes that were not expressed at all were mapped to X = 0, and the median of all genes in a replicate was mapped to *X* = 10. The mean and standard deviation (sd) of expression levels over replicates was computed for each gene-tissue pair to generate a distribution of 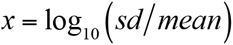 for human and mouse data respectively (Fig S1). Based on these distributions cutoffs *x_h_*and *x_m_* were established and the gene-tissue pair was excluded from further analysis if *x* was greater *x_h_* ~ -0.4 and *x_m_* ~ -0.6 for human and mouse data respectively. For the gene-tissue pairs that passed this check, the expression of a gene in a tissue was defined as the mean over replicates. If a gene had to be excluded in many tissues, then that gene was excluded altogether. Sixty-six ribosomal proteins in 110 tissues in humans (S7 Table), and 42 ribosomal proteins in 18 tissues in mice (S8 Table), passed this filter. The R package *pvclust* was used to perform bootstrapping [36] of hierarchical clustering of ribosomal proteins and tissues in mouse and humans. Similar filtering was performed for GTEx data. GTEx data consists of RNAseq data of 54 tissues from 544 donors amounting to a total of 8,555 samples. Gene-tissue pair with *x_g_* > 0.1 were excluded from the data. Seventy-nine ribosomal proteins in 54 tissues passed this filter (S9 Table).

## RESULTS

### Phenotypic variability of ribosomal proteins in yeast

In all organisms, the ribosome is a ribonucleoprotein complex composed of two subunits each with an RNA core and large number of ribosomal proteins. In eukaryotes, the 60S large subunit consists of 46 proteins, and the 40S small subunit consists of 33 proteins [37]. In yeast that has undergone whole genome duplication [38], most of the ribosomal proteins (paralogs) are duplicated, as a result of which it contains 137 ribosomal proteins, of which 107 are non-essential [22].

Deletion collection in yeast allows testing of phenotypic effect of deletions of non-essential genes in yeast in diverse environments. In order to identify genes which show maximum phenotypic variability across environments, we reanalyzed deletion phenotypes for 4,769 single gene deletions grown in 293 diverse environments using a previously published dataset [22] (S1 Table) for all genes, including ribosomal proteins. The surprising observation was that across all yeast genes, deletions of ribosomal proteins had the highest differential effect on growth in different environments i.e., no effect in some environments and strong effect in others. Among the 191 genes with variance σ ^2^ > 0.8 across the 293 environments, components of the ribosome were significantly enriched (21/191, *P* < 0.01, S10 Table, Fig S2). Fourteen out of these 21 genes belonged to the large ribosomal subunit. These 21 genes contain only one paralog of the ribosomal proteins, either A or B, indicating a possible but small differential role of paralogs in responding to environmental heterogeneity. Only ribosomal protein *RPL34* was an exception to this where both paralogs - *RPL34A* and *RPL34B* showed high phenotypic variance.

To test whether this phenotypic variability was a non-specific cellular effect or was specific to the ribosome, we compared phenotype variability in growth for deletions of genes in 90 different pathways and protein complexes across 293 stress conditions versus growth in rich media (YPD). These 90 pathways and complexes were defined using both a biased (Wikipathways) [26] and an unbiased (SGA clustering) [23] approaches (see Methods) and included signaling pathways such as the MAPK and Ras/PKA pathway, protein complexes such as the proteasome and ribosome, cellular processes such as chromatin remodeling complexes and vesicular transport machinery, etc. (S2 Table). Differences in variance of a pathway or a complex between stress and YPD indicate variable roles of its constituents in different conditions. High correlation of constituents of a pathway between stress and YPD would indicate that independent of the essentiality of the pathway; different constituents have similar functions in both conditions. Moreover, a higher variance in YPD compared to stress would indicate that the constituents of the pathway show a more diverse response in YPD but show similar phenotype in stress (Fig 1A). Such a co-ordination of stress specific genes has previously been observed [39] in multiple stresses where the whole pathway is essential to respond to the stress. On the other hand, higher variance of the pathway in stress compared to YPD would indicate that different components of the pathway have differential roles in stress and therefore function in a different manner than in YPD (Fig 1B). Deletions of constituent proteins in 13 pathways showed a significant difference in variance in 3 or more stress environments compared to YPD (*P* < 0.01 by Brown Forsythe test, S2 Table, Fig S2), with the higher variance in YPD in most cases, showing that the pathway was essential in stress. Additionally, constituent of these pathways showed high correlation of phenotype across YPD and stress indicating that the functional hierarchy of the genes was conserved (functional homogeneity), but the phenotypic contribution of the module increased during stress, reducing phenotypic variance. In contrast, for the cytoplasmic ribosomal proteins, there was a significantly higher variance in 28 stress environments compared to YPD (Fig S2, S2 Table), suggesting that ribosomal proteins are differentially used in the stress condition. Unlike other pathways, poor correlation was observed between phenotype of ribosomal proteins across YPD and stress, indicating overall functional heterogeneity in stress and YPD (Fig 1B). These results independently show that among different pathways, deletions of genes in the ribosomal pathway has the greatest effect on growth in stress versus rich media, suggesting a unique property of the ribosomal genes that they are the most variable proteins in the cell when comparing diverse environments. A heatmap of growth for single deletions of the 68 ribosomal proteins with consistent replicate data in 25 stress conditions and YPD (Fig 1C) reinforces the above results and shows that a number of these ribosomal proteins have high phenotypic variability, i.e. that they are required for growth in some environments but expendable in others.

**Fig 1:**
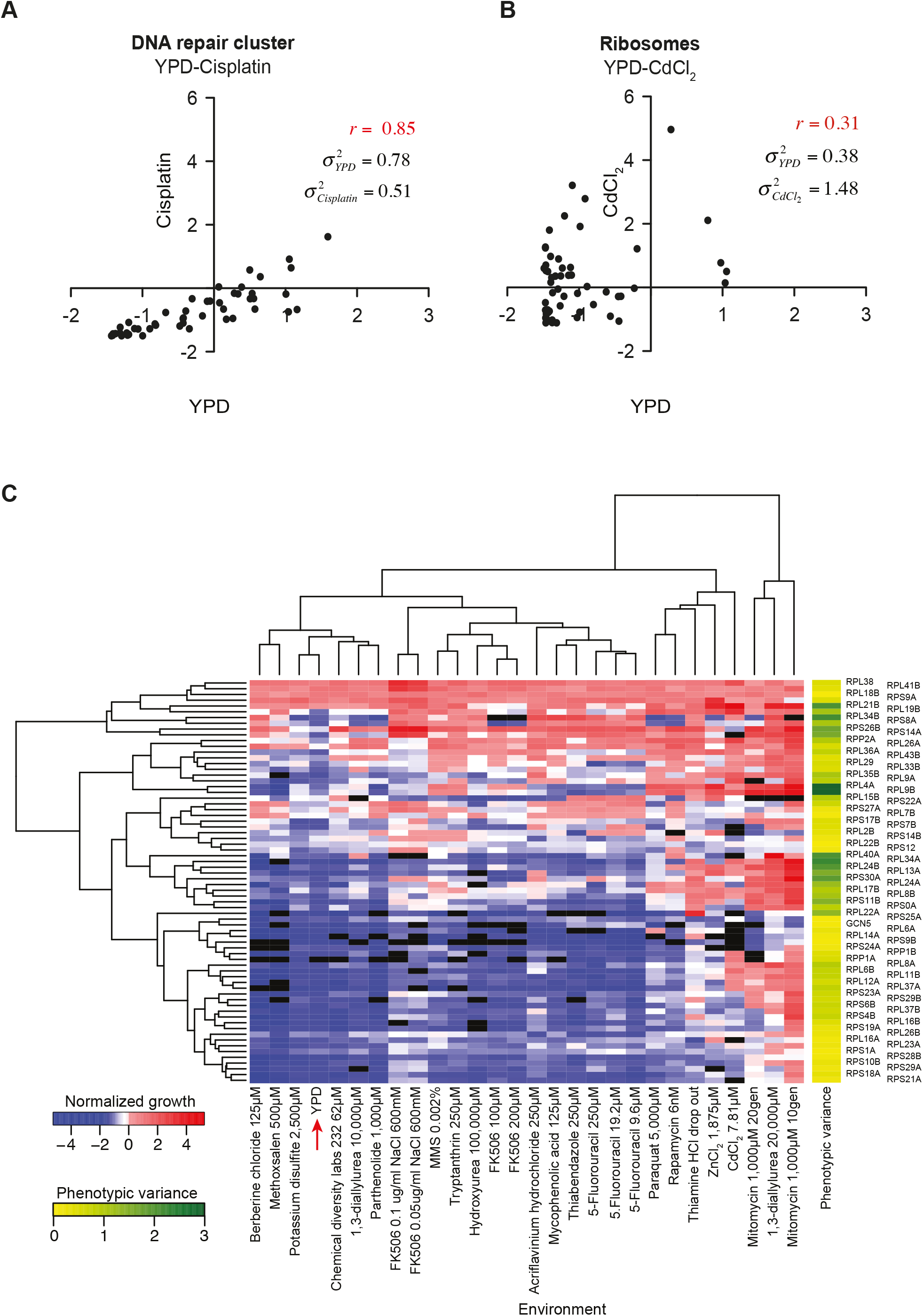
Phenotypic variability of yeast ribosomal proteins. (A) Growth of deletions of DNA repair cluster genes (black dots) in rich medium YPD (x-axis) versus a DNA damaging agent, cisplatin (y-axis). (B) Growth of deletions of ribosomal proteins (black dots) in rich medium, YPD (x-axis) versus an oxidizing stress, Cadmium chloride (CdCl_2_) (y-axis). (C) Hierarchical clustering heat map of normalized growth of yeast strains for 68 single deletions of ribosomal proteins in 26 environments. The red arrow indicates YPD (rich medium).

We next asked whether these ribosomal proteins have different phenotypic profiles across environments i.e., whether they work independently, or whether they form modules, whose constituents show coordinated regulation across different environments. A clustering analysis of the Pearson correlation of these ribosomal proteins across environments showed functional modularity (Fig 2A) in the form of three distinct clusters (S11 Table). Ribosomal proteins in Cluster 1 were both highly correlated, enriched in large subunit proteins and had high phenotypic diversity across environments. On the other hand, proteins in Cluster 2, although highly correlated and important for growth across most environments, were enriched in small subunit and pre-ribosomal components (important for ribosomal assembly), which explains the constitutive growth defect when these proteins were deleted (S10 Table). Proteins in Cluster 3, however, showed low correlation amongst themselves and were important in different environments. The conclusion that emerges from this analysis is that subsets of ribosomal proteins in Cluster 1 act together in diverse environments, whereas proteins in Cluster 2 act together in most environments. Proteins in Cluster 3 on the other hand, seem to play specialized roles in specific environments.

**Fig 2:**
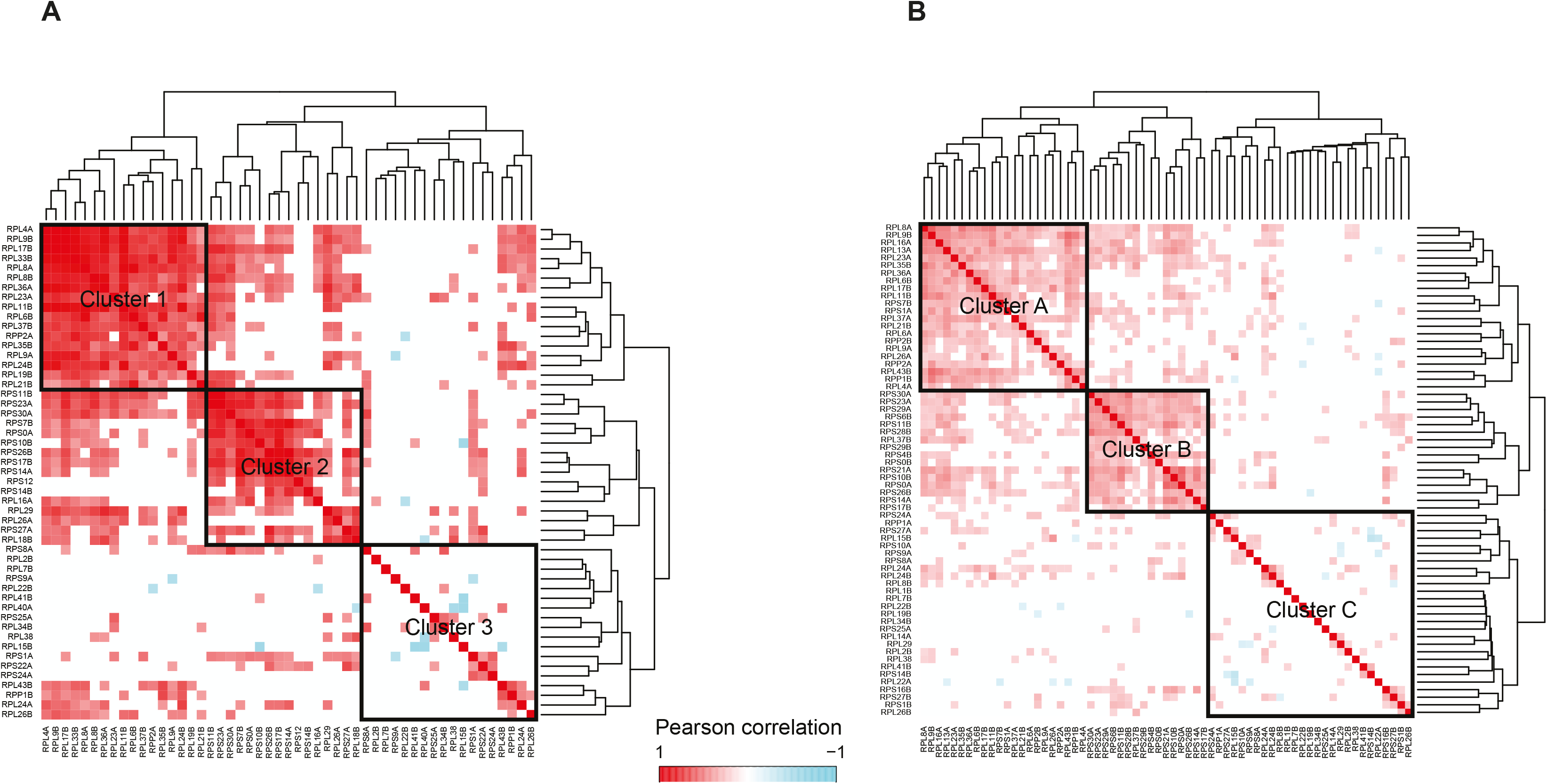
Differential modules of ribosomal proteins in yeast. Hierarchical clustering of Pearson correlations among ribosomal proteins for (B) normalized growth from single deletion of ribosomal proteins in 26 environments, (C) double deletion interactions with 121 genes. Ribosomal proteins within each cluster are significantly correlated with *P* < 0.05 for each pair.

These results strongly suggest that the yeast ribosomal proteins do not function in a uniform manner when the environment is varied. While this environmental variability of deletions of ribosomal proteins has previously been observed, we have identified a novel underlying modularity among these ribosomal proteins, potentially to optimize yeast growth in different environments. *This is the main finding of the present work that distinguishes it from previous studies.* In summary, whereas a core set of ribosomal proteins are important in all environments, different combinations of a subset of variable ribosomal proteins are functional in different environments to optimize growth.

To test whether the deletion phenotype of ribosomal proteins is conserved among yeast strains, we compared growth of ribosomal protein deletions in a soil isolate, SK1, in an oxidative stress (Fig S3). While different ribosomal protein deletions show diverse phenotypic defects, indicating differential use of ribosomal proteins in SK1, the identity of the variable ribosomal proteins was different among the two strains, SK1 and S288c. This indicates that different strains functionally employ ribosomal proteins in different ways, potentially as a result of having adapted in different ways to strain specific selection pressures.

### Interactions among ribosomal proteins and genes in cellular pathways

If this observed phenotypic modularity of ribosomal proteins is indeed real, then it should be reflected in their genetic interactions with both upstream and downstream pathways. This crosstalk between ribosomal proteins and genes in other pathways was captured by studying the genome-wide gene-gene interaction SGA dataset [23], to identify positive and negative genetic interactors of the ribosomal proteins. A positive or negative interaction is one where the double deletion is respectively better or worse for growth than the sum of the single deletions. A positive interaction indicates that the genes are in the same pathway, while a negative interaction indicates compensatory pathways [26].

The interactions of a diverse set of genes with 65 ribosomal proteins, 57 of which overlapped with the ribosomal proteins in the deletion phenotype analysis, was analyzed using the SGA dataset [23]. While multiple genes showed double deletion genetic interactions with the ribosomal proteins, our aim was to identify genetic interactors that are common to the ribosome and not to a single ribosomal protein. Hence, only those genes that showed a significant interaction (*P* < 0.05) with at least 10 of the 65 ribosomal proteins were considered for further analysis. This identified a total of 121 genes, 23 ribosomal and 98 non-ribosomal, which had a significantly positive or negative interaction with at least 10 ribosomal proteins (modified F-test, *P* < 0.05, S3 Table). Although the 65 ribosomal proteins had many interactions with other genes, only 12 ribosomal proteins interacted positively among themselves (i.e., were in the same pathway). The rest interacted either positively or negatively with non-ribosomal proteins. The ribosomal proteins form a separate cluster in the yeast gene-gene interaction dataset [23] due to an enrichment of interactions within the complex. Our results show that this enrichment is the result of only 12 ribosomal proteins (enriched in small ribosomal subunit); the other ribosomal proteins show high interactions with different cellular pathways.

If the ribosomal proteins indeed function in a coordinated and modular manner to regulate growth across environments, then this modularity should also be evident in their genetic interactions. A clustering analysis of the pairwise correlations of the positive or negative genetic interactions of the 65 ribosomal proteins (Fig 2B) identified three clusters, which had a highly significant overlap with the previous clustering based on phenotypic profiling (90% overlap between Clusters 1 and A and 89% between clusters 2 and B, Fisher Exact test, *P* = 0.006, S4 Table). Ribosomal proteins in Cluster A interacted mainly with genes involved in mRNA processing, whereas those in Cluster B interacted with other ribosomal proteins. Ribosomal proteins in Cluster C interacted with genes involved in diverse pathways (S10 Table). This strong overlap of corresponding clusters identified independently through phenotype association (Clusters 1, 2, 3) and double deletion analysis (Clusters A, B, C) further reinforces our claim of the modularity of ribosomal proteins wherein subsets of ribosomal proteins function in a coordinated manner and interact in diverse ways amongst themselves and with other non-ribosomal pathways.

We observe that despite high sequence similarity, sometimes differing in only a few bases, ribosomal proteins and their paralogs show diverse genetic interactions. Only 4 out of 15 paralog ribosomal proteins show similar genetic interactions i.e., fall in the same cluster. These are *RPL6A* and *RPL6B*, and *RPL9A* and *RPL9B* in Cluster A and *RPS0A* and *RPS0B*, and *RPS29A* and *RPS29B* in Cluster B (S11 Table). All the remaining paralogs fall into separate clusters or are in Cluster C, indicating that they have differential genetic interactions. Thus, in spite of sequence similarity amongst paralogs, the regulation of these modules in yeast seems to have evolved since the duplication event to create novel functions for these paralogs.

Positive non-ribosomal interactors of the ribosomal proteins were enriched in chromatin regulators and remodelers (S10 Table). Under the prevailing unitary ribosome hypothesis, this association is believed to result from a general control of cellular proliferation by epigenetic regulators [40]. To test this, we experimentally studied the observed positive interaction between the chromatin histone deacetylase *GCN5* and the ribosomal proteins *RPL11B* (Cluster A), *RPL6B* (Cluster B), *RPL38* (Cluster C) and *RPL26B* (Cluster C). *GCN5* regulates transcription of these ribosomal proteins, presumably to control cellular proliferation [40]. If this were true, then deletion of *gcn5Δ* with deletion of any of these ribosomal proteins should have the same growth phenotype as single deletion of *gcn5Δ*. Since ribosomal proteins showed diverse phenotypes in oxidative stresses (Fig 3), we tested the premise of these genetic interactions by phenotyping double deletions of *GCN5* and ribosomal proteins in YPD, and oxidative stresses menadione and CdCl_2_. While *gcn5Δ rpl38Δ* behaved the same as *gcn5Δ* in menadione, indicating that *GCN5* controls cellular proliferation using *RPL38,* double deletion *gcn5Δ rps6bΔ* resulted in a 20-fold reduced growth compared to either of the single deletions, indicating parallel or independent roles of both the genes in growth. Furthermore, deletion of the remaining ribosomal proteins (*rpl11bΔ* and *rpl26bΔ*) rescued the growth phenotype of *gcn5Δ* (Fig 3). This observed antagonistic effect suggests that a more likely scenario is that these ribosomal proteins have a direct functional effect on pathways that are lost as a result of *gcn5Δ* deletion. This varied environmental dependence of genetic interactions of ribosomal protein and *GCN5* indicates that some epigenetic regulators and signaling pathways that interact with the ribosome employ the flexibility of the ribosome to mediate phenotypic choice in diverse environments.

**Fig 3:**
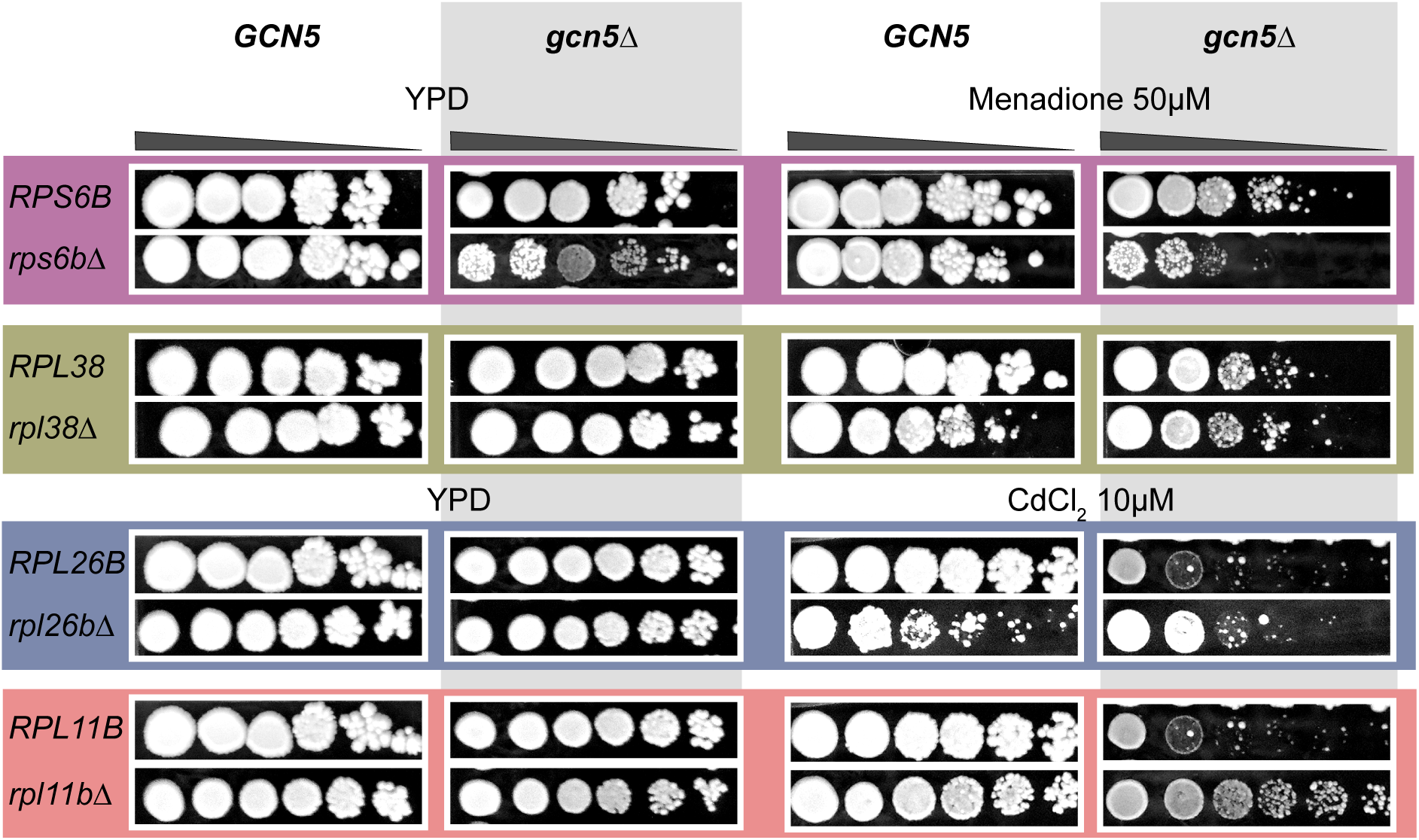
Genetic interactions of ribosomal proteins with *GCN5*. 10-fold spot dilutions series (starting with 10^8^ cells/ml) of wild type, single and double deletions of *GCN5* with *RPS6B*, *RPL38*, *RPS26B* and *RPL11B* phenotyped in YPD (rich medium) and an oxidative stress, Cadmium chloride (CdCl_2_, 10µM).

### Differential evolution of ribosomal proteins

If different subsets of ribosomal proteins are under selection for adaptation to different environmental conditions, there should be a signature of this effect in yeast strains adapted to different environments. The yeast SGRP population [32] consists of 38 *S. cerevisiae* strains isolated from diverse ecological and geographical niches. To detect potential footprints of evolutionary selection, we compared the nucleotide diversity of ribosomal proteins and a control set of housekeeping genes in the SGRP population in their coding and 5’UTR sequences (S5 Table). While the coding sequence nucleotide diversity was similar for both sets, the diversity in the 5’UTR regions within ribosomal proteins was twice that of the control genes (*P* < 0.005, Fig S4A, S5 Table). Furthermore, in the YEASTRACT database [35], this variability altered the predicted transcription factor binding motifs on ribosomal proteins in diverse strains (S7 Table). Thus, while ribosomal proteins among the SGRP strains have similar coding sequences, their promoter regions have been significantly altered, presumably to adapt to different ecological niches.

Coding and 5’UTR regions of ribosomal proteins in the three clusters in Fig 1C were compared using normalized Shannon entropy (S5 Table). All three clusters showed a significant difference in entropy between 5’UTR and coding regions (Fig S4B). Cluster A showed significantly high entropy (diversity) for both 5’UTR and coding regions of ribosomal proteins compared to Clusters B and C (Fig S4B). This differential variability shows that the ribosomal proteins in the clusters are evolving at different rates across the SGRP population.

Using the YEASTRACT database, we find that whereas most transcription factors (TFs) bind to ribosomal proteins in all three clusters, some are cluster specific. Transcription Factors regulating ribosomal proteins in Cluster A are enriched in the histone deacetylase complex while those that bind to ribosomal proteins in Cluster C are enriched in the HIR (Histone Regulatory) complex (Fig S5, S6 Table). This could be a possible explanation for the different rates of evolution of the 5’UTR regions of the proteins in these three clusters.

### Modular ribosomal proteins in higher eukaryotes

Our above results establish modularity in both phenotype and genetic interactions of ribosomal proteins in yeast. It can be argued that this is merely a unique feature of the yeast ribosome, presumably because of the whole genome duplication event, which might have allowed differential adaptation of duplicated ribosomal proteins. To understand whether the ribosomal modularity observed in yeast extends to higher eukaryotes, we investigated expression levels of ribosomal proteins in mice and humans, which have a single copy of most ribosomal proteins.

In complex eukaryotes, the analog of adaptation of unicellular organisms like yeast to different environments is adaptation to different cellular and tissue microenvironments. We therefore expect that if our thesis of the modularity of ribosomal proteins is valid beyond single celled eukaryotes, ribosomal proteins should be differentially used across cell types and tissues in mice and humans. In multicellular systems like humans and mice, ribosomal proteins are present in a single copy, whose deletion results in both cellular and organismal lethality. Our hypothesis for complex eukaryotes would then be that ribosomal proteins are expressed at significantly different levels in different tissue microenvironments in mice and humans. We tested this hypothesis by comparing the expression levels of ribosomal proteins in diverse cell types and tissues in human and mouse samples using RNASeq data for mRNA transcript levels from the ENCODE and the GTEx projects.

A total of 66 ribosomal proteins with consistent transcript expression levels across replicates in the ENCODE data were identified in 110 cell types and tissues in humans (see Methods). As previously observed [41], we found that while the majority of ribosomal protein transcripts are highly expressed across diverse tissues and cell types, a few showed low expression levels throughout. It has been shown that ribosomal proteins can show differential expression levels based on the proliferation or turnover rate of the cell type. To normalize such global differences, each ribosomal protein within each tissue was assigned a rank based on its expression level (rank 1 for lowest expression and rank 66 for highest expression, S7 Table). Despite being a part of the ribosome, it is known that not all ribosomal proteins are equally expressed in a given tissue. In complex eukaryotes, just as in yeast, some of the ribosomal proteins are involved in ribosomal assembly. However, since this is true for all tissues and cell types, their rank normalized expression levels should be consistent across tissues.

Hierarchical clustering of all ribosomal proteins expression ranks across all tissues resulted in a single highly correlated cluster (Fig S6). However, our results from yeast show that while some ribosomal proteins are essential and behave similarly across environments, others show high variability. To identify these highly variable ribosomal proteins, the 66 ribosomal proteins were filtered based on their ranks. In each tissue, the expression level ranks of the proteins were stratified into 4 classes: class I (rank 1-17), II (17-34), III (35-51) and IV (51-66) (see Methods). This showed that across tissues, 46 out of 66 ribosomal proteins were classified into the same or adjacent classes, while the remaining 20 were classified into 3 classes for 11 or more tissues types per class (S7 Table). These 20 were termed as variable ribosomal proteins and analyzed further. Note that had the rank assignments merely amplified small differences in expression levels (noise) for a given protein across tissues, such stratification would not have been observed. Instead, we would have seen a random assignment of ranks across tissues, which is not what was observed.

The 20 variable ribosomal proteins spanned mostly classes II, III and IV i.e., their transcripts were both highly expressed and highly variable across tissues, thereby eliminating technical noise as the cause of the observed variability in ranks (S7 Table). Hierarchical clustering of the ranks of these 20 proteins across cell types and tissues showed that these proteins assort into distinct groups (Fig 4A). An identical clustering can also be observed in a heat map of Pearson rank correlations (Fig 4B, correlation *P* < 0.05). In the hierarchical clustering, distinct sets of ribosomal proteins were associated with two discrete clusters of epithelial cells, a cluster of the nervous tissue (tissue from different sections of the brain and spinal cord) and a cluster of human cell lines (Fig 4A). We note that cell lines cluster separately, indicating that similar to modification of their signaling pathways [42], expression patterns of ribosomal proteins are also rewired in these cell lines compared to other human cells and tissues. To investigate this tissue specific ribosomal modularity further, we separately analyzed the 20 ribosomal proteins in the clusters associated with epithelial and nervous tissues. This again showed that the nervous tissue cluster is quite distinct in its use of variable ribosomal proteins compared to epithelial cells (Fig 5A, 5B). Even though the nervous tissues are known to have a reduced expression of ribosomal proteins compared to other more proliferating tissues, our results show that the ribosomal protein module in the nervous tissue is distinct from that in the epithelial cells. These results show that, analogous to yeast, ribosomes in humans show expression modularity across tissues.

**Fig 4:**
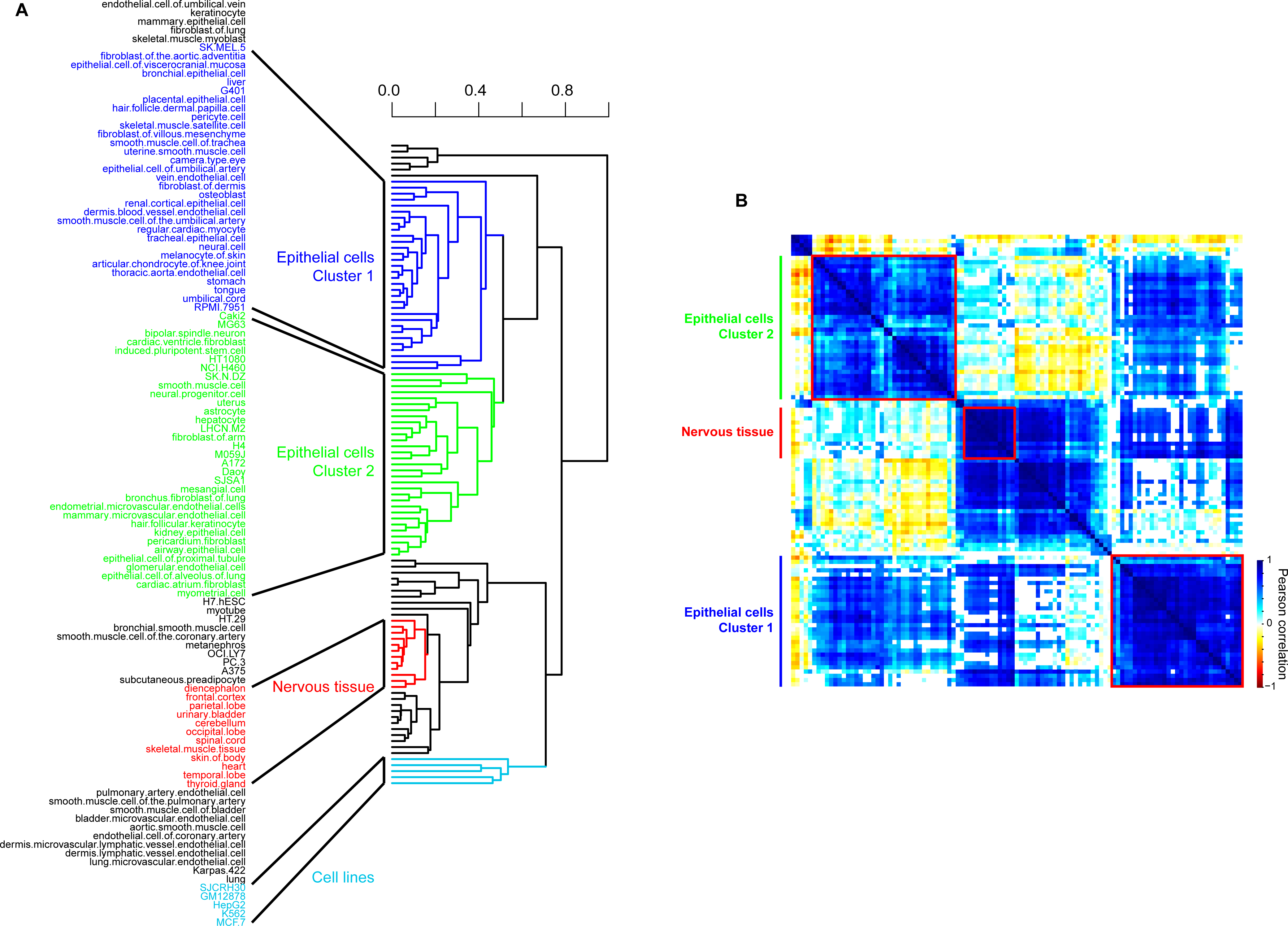
Clustering of human cell types and tissues based on rank order expression of variable ribosomal proteins in ENCODE data. (A) A hierarchical clustered tree of 110 human cell types and tissues (1,000 bootstraps) based on rank order expression of 20 ribosomal proteins in ENCODE data. (B) Pearson correlation (*P* < 0.05) heatmap based on rank order expression of 20 variable ribosomal proteins results in distinct clusters and sub-clusters.

**Fig 5:**
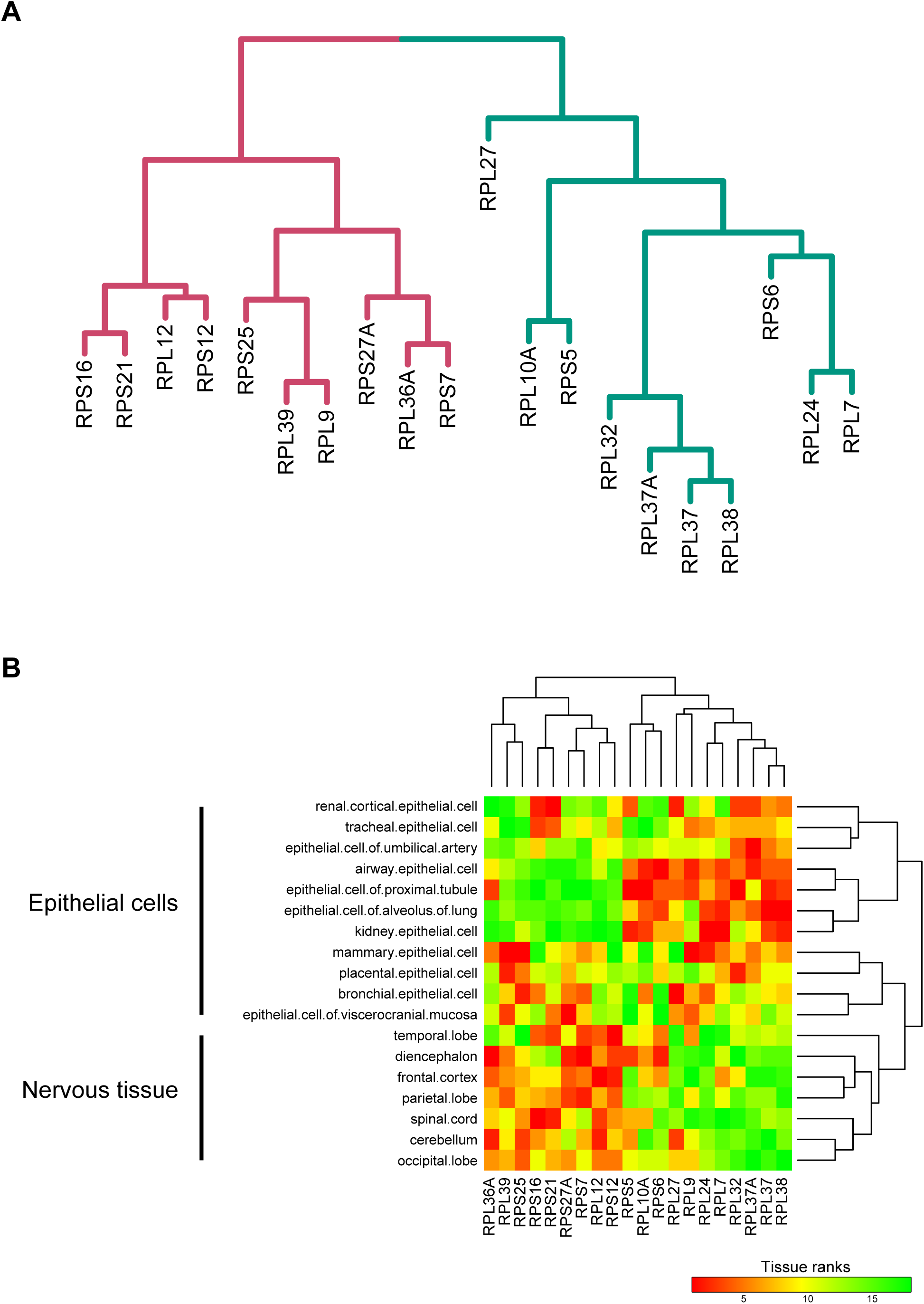
Modules within the 20 variable ribosomal proteins in human tissues. (A) The modular organization derived from hierarchical clustering of ribosomal proteins rank order expression in 110 cell types and tissues (1,000 bootstraps). (B) Heatmap showing different modules of 20 variable ribosomal proteins active in epithelial cells and the nervous tissue.

While the different cell types separated into different clusters based on the expression of variable ribosomal proteins in ENCODE data, they did not show a significant tissue bias except for nervous tissue. For example, the two clusters of epithelial cells did not segregate on the basis of organ of origin, suggesting that modularity of ribosomal proteins plays a role at the resolution of cell types instead of whole organs. To validate our result of modularity of ribosomal proteins in humans and to understand whether this modularity was a property of cell types or is also visible in bulk tissue, we compared the expression patterns of ribosomal proteins using the GTEx dataset. The GTEx data consists of RNAseq analysis of 54 different tissues (S9 Table). We extracted expression values of ribosomal proteins from these samples and performed the same analysis as for the ENCODE data (see Methods). Seventy nine ribosomal proteins passed our filtering criteria and were classified into ranks ranging from 1 for the least expression and 79 for the highest expression in each tissue. These were further stratified into 4 classes: class I (rank 1-20), II (21-40), III (41-60) and IV (61-79). We found that ribosomal proteins from the GTEx data showed less variability in classes across tissues compared to the ENCODE data (S9 Table). Consequently, ribosomal proteins that fell into two or more classes with at least 10 tissues per class were identified as variable ribosomal proteins. A total of 18 ribosomal proteins were identified to be variable, of which 7 were the same as those identified in the ENCODE data, showing a significant overlap (Fisher’s Exact test, *P* < 0.1) between variable ribosomal proteins identified using ENCODE and GTEx data (Fig 6A). The variable ribosomal proteins in the GTEx data separated into two modules (Pearson correlation, *P* < 0.01, Fig 6A). Similar to the ENCODE data, the nervous tissues (brain and spinal cord) formed a separate cluster, validating our previous observation that a different module of ribosomal proteins is used in the nervous system compared to other tissues (Fig 6B). However, bulk tissues did not cluster separately on the basis of variable ribosomal proteins (Fig 6B). The lack of modularity of ribosomal proteins at the level of tissues indicates that cell specific differences in ribosomal protein expression levels are lost when dealing with data from bulk tissue, because cell specific identity is lost in the GTEx data.

**Fig 6:**
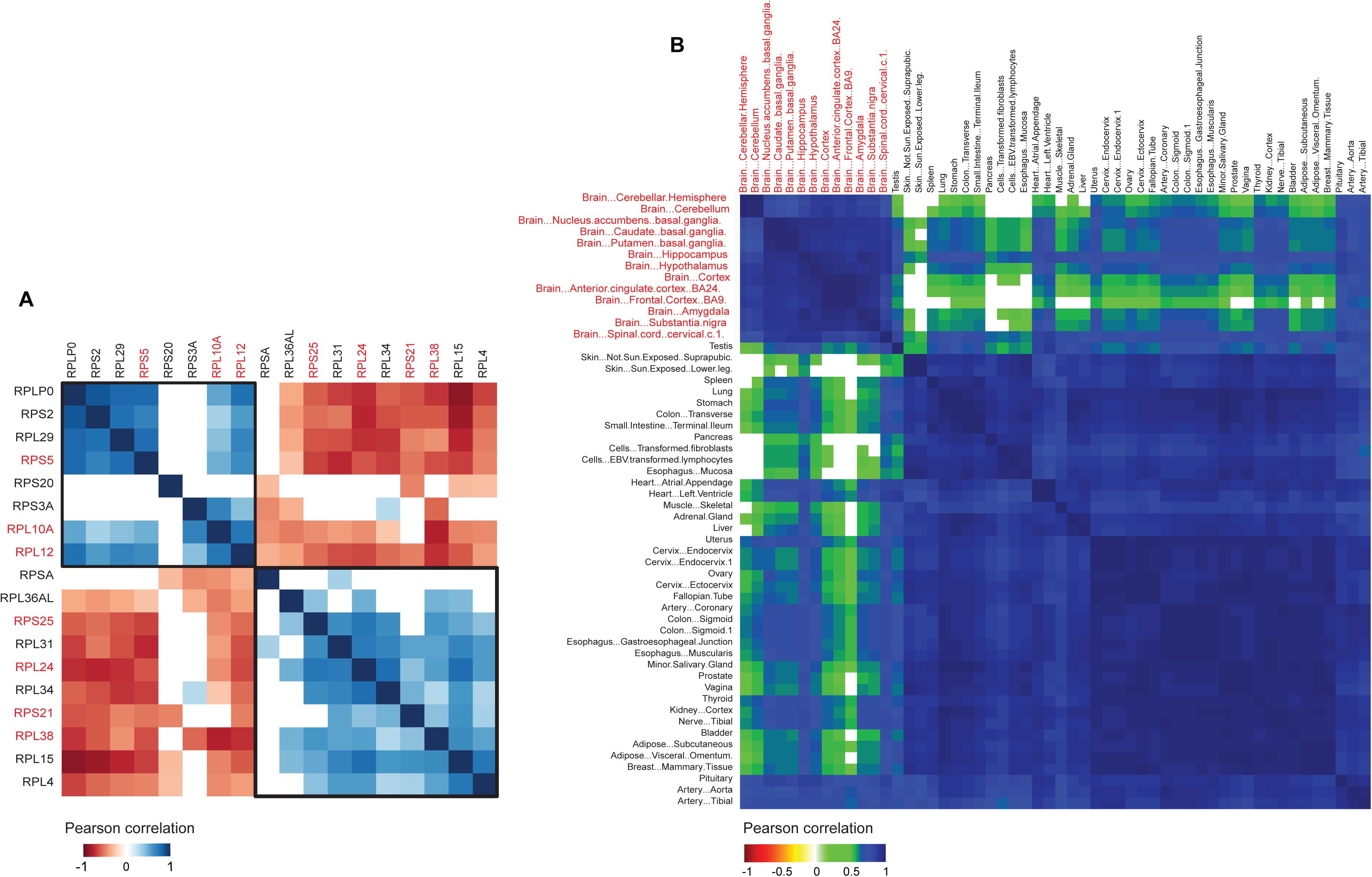
Clustering of human tissues and variable ribosomal proteins in GTEx data. (A) Pearson correlation (*P* < 0.01) heatmap of 18 variable ribosomal proteins based on their rank order expression in 54 different tissues in GTEx data. Ribosomal proteins marked in red are also variable in ENCODE data. (B) Pearson correlation (*P* < 0.01) heatmap of 54 different tissues based on rank order expression of 18 variable ribosomal proteins. Nervous tissue (brain and spinal cord) form a distinct cluster and are marked red.

A similar analysis was carried out for expression levels of ribosomal proteins in various tissues in mice from the ENCODE data. We note that the signal to noise ratio in the mouse data in ENCODE was significantly higher than in the human data. Consequently, only 42 ribosomal proteins in 18 different tissues passed our filtering criteria and were stratified into four classes (S8 Table for details of classes). As in the human data, while the majority of mice ribosomal proteins showed high expression and invariant classification, 14 of these 42 ribosomal proteins were as variable across tissues (spanning 3 classes in more than 2 tissues). As in ENCODE and GTEx data from humans, mouse brain tissues also formed a unique cluster, indicating that nervous tissue in general uses a distinct module of ribosomal proteins compared to other tissues (Fig S7A, 7B).

We identified *RPL38* as a variable protein in ENCODE and GTEx data in humans as well as ENCODE data in mice (Fig 5, 6, S7B). *RPL38* is the most extensively studied ribosomal protein associated with ribosomal heterogeneity. This heterogeneity is due to its specialized translation of only the *hox* mRNA, without affecting translation of other mRNA [19]. Identification of *RPL38* as a variable ribosomal protein in our study serves as an independent validation of our analyses. While different sets of ribosomal proteins were found to be variable in mice and humans, 6 out of 18 ribosomal proteins variable in mice were also variable in humans (4 in ENCODE data and 5 in GTEx data, Fig S7B). Furthermore, 4 of these 6 conserved variable ribosomal proteins fell in one cluster in mouse indicating a partial conservation of variability of ribosomal proteins across the two species (Fig S7B).

## DISCUSSION

Our study provides several arguments and multiple evidences for the existence of modularity of ribosomal proteins across eukaryota, presumably to facilitate optimized translation efficiency in different environments. We show that, at least in yeast, we see evidence that the 5’UTRs of ribosomal proteins that form the modules seem to be under selection pressure, which suggests that they play a role in evolutionary adaptation. We interpret our results as evidence for a hitherto unrecognized ribosomal code, wherein specific ribosomal proteins are used in an environment specific manner in yeast and in cell and tissue specific ways in mice and humans. *The existence of such a dynamic modularity of ribosomal proteins is the main finding of this paper.* The mechanisms that regulate these modules remain to be elucidated and are outside the scope of this paper.

Our study also uncovered some general, conserved properties of ribosomal proteins. We find that a subset of variable ribosomal proteins contribute to the plasticity of the ribosome by functioning independently or in concert across different environments by forming *modules*, defined as sets of proteins functioning in a coordinated manner. This modularity indicates that they have been optimized over the course of evolution based on the need for functional adaptation. We note that only a subset of ribosomal proteins vary among cell types and tissues, with the core ribosome remaining unaffected. Hence cell type specific structural changes resulting from such variation may be difficult to detect.

Our findings would argue that, in spite of high sequence conservation [43], the inability of human ribosomal genes to substitute for yeast ribosomal genes [44] is probably because of species specific functioning of ribosomal modules. This, along with differential expression variability of ribosomal proteins in mice and humans, indicates that each species optimizes the composition of its ribosome to adapt to species specific selection pressures, not by substantially altering the sequence of the ribosomal proteins but by regulating their expression in an environment dependent manner using mechanisms yet to be discovered.

Our results from two independent expression datasets (ENCODE and GTEx) show that the nervous tissues use a unique ribosomal code compared to the rest of the tissues in both mice and humans. While an overall reduced expression of ribosomal proteins in the brain has been observed previously, it has been attributed to the reduced proliferation of nervous cell types. Here, we show that along with a reduced expression, a unique composition of ribosomal proteins is utilized by nervous tissue. These may play a role in the fundamental physiological differences observed between the brain and the rest of the body.

In mice and humans, recently evolved paralogs *RPL27L*, *RPL22L1*, *RPL7L1*, and *RPL39L*, showed poor but highly tissue specific expression compared to core ribosomal proteins. This suggests that there is an ongoing process of adaptation driving modular ribosomes, with recent paralog proteins still evolving in response to selection pressures on them and on other, more ancient ribosomal proteins and pathways.

Our results show that ribosome modularity is a dynamic, evolving process which seems to be involved in the evolution of species specific ribosomal proteins, the diversification of their sequences and functions and the creation of novel, species specific ribosomal proteins [45,46] leading to diverse phenotypic adaptations.

## SUPPLEMENTARY INFORMATION

**Fig S1: Distribution of signal to noise in human and mouse data from ENCODE.**

ENCODE data was normalized by median subtraction per array and then log transformed (see Methods). The mean and standard deviation (sd) over replicates was computed to obtain 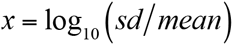. The distribution of *x* for human (A) and mouse (B) ENCODE data and (C) human GTEx data was used to determine the cutoffs *x_h_*, *x_m_*and *x_g_* for reliability of the ENCODE data for human, mouse and GTEx data for humans, respectively. Gene-tissue pairs for which *x_h_*> -0.4, *x_m_*> -0.6 and *x_g_*> 0.1 were excluded from the analysis.

**Fig S2: Phenotypic variation of yeast ribosomal proteins.**

(A) Distribution of variance of normalized growth of all non-essential genes in yeast (4,769) in 293 different environments. The genes are on the x-axis, arranged in increasing order of the variance of normalized growth in 293 environments (y-axis) due to their deletion. The 191 genes to the right of the red line have variance greater than 0.8. (B) Stacked histogram showing the number of cases when the variance of growth from deletion of genes in various pathways was greater or smaller in stress compared to YPD. Gene deletions in the cytoplasmic ribosome and mitochondrial tRNA synthesis pathways had the highest variance in stress compared to YPD. However in mitochondrial tRNA synthesis pathway genes, the variance is greater or smaller in stress compared to YPD for equal numbers of genes. Only in the ribosomal pathway is the variance in stress conditions greater than YPD for all genes. (C) This figure shows the same data as in B but with the genes stratified into clusters based on their double deletion interactions (see Methods). The ribosomal cluster has the highest variance in stress compared to YPD.

**Fig S3: Phenotype of ribosomal protein deletions in SK1 strain**

Ten-fold spot dilutions series (starting with 10^8^ cells/ml) of wild type and ribosomal protein deletion strains of SK1 background phenotyped in rich medium YPD and an oxidative stress, Cadmium chloride (CdCl_2_ 500 µM)

**Fig S4: Nucleotide diversity of ribosomal proteins across the SGRP strains:**

(A) Nucleotide diversity of coding and promoter sequences of ribosomal proteins and a control set of genes using Tukey’s multiple comparisons’ test (*P* < 0.05). The bars with the same letter code do not differ significantly.

(B) Normalized Shannon Entropy of coding region and 5’UTRs of Cluster A, B and C from Fig 2B. Bars with the same letter code do not differ significantly (Tukey’s multiple comparisons’ test, *P* < 0.05). The figure shows that: (i) The 5’UTR regions of the ribosomal protein sequences are most variable compared to their coding region as well as the 5’UTR and coding regions of the control set of genes; (ii) The 5’UTRs of all the ribosomal proteins in the three clusters are significantly more variable (*P*<0.01) than their coding regions; (iii). Proteins in Cluster A have significantly more variable coding regions than clusters B and C (*P* < 0.05 and *P* < 0.01 respectively); (iv). Proteins in Cluster A have significantly more variable 5’UTR than Clusters B and C (*P* < 0.05 and *P* < 0.01 respectively).

**Fig S5: Networks of transcription factors that bind uniquely to ribosomal proteins in Cluster A, B and C (see Fig 2B).**

These network clusters were identified using the STRING database (http://string-db.org). The thickness of blue lines connecting two transcription factors indicates the strength of experimental evidence for their interaction. Gene enrichment (*P* < 0.001) of transcription factors in Cluster A is for Histone Deacetylase Complex and in Cluster C for the HIR Complex.

**Fig S6: Pearson correlation heatmap (*P* < 0.05) of 110 human cell types and tissues based on rank order expression of 66 ribosomal proteins in ENCODE data.**

**Fig S7: Variable ribosomal proteins across tissues in mouse**

(A) Hierarchical clustering of 18 tissues in mice based on expression rank orders of 14 variable ribosomal proteins (1,000 bootstraps).

(B) Hierarchical clustering of the 14 variable ribosomal proteins based on their expression rank orders in 18 tissues in mice (1,000 bootstraps). The ribosomal proteins form three distinct clusters indicated in different colors. The red arrows indicate ribosomal proteins that are also variable in human ENCODE and GTEx data.

**S1 Table:** Yeast deletion collection phenotyped in 293 environments.

**S2 Table:** Variance analysis of various pathways and genetic

**S3 Table:** Double deletion genetic interactions of the 65 ribosomal proteins with 121 ribosomal and non-ribosomal proteins.

**S4 Table:** Strains used in this study and Primer sequences

**S5 Table:** Nucleotide diversity and Shannon Entropy comparisons of ribosomal proteins in Clusters A, B and C.

**S6 Table:** Transcription Factors binding to ribosomal proteins in Clusters A, B and C downloaded from YEASTRACT

**S7 Table:** Expression ranks and classes of 66 ribosomal proteins in 110 human cell types and tissues downloaded from ENCODE

**S8 Table:** Expression ranks and classes of 42 ribosomal proteins in 18 mouse tissues downloaded from ENCODE

**S9 Table:** Expression ranks and classes of 79 ribosomal proteins in 54 human tissues downloaded from GTEx portal.

**S10 Table:** Gene Ontology Enrichment for various groups and clusters

**S11 Table:** ribosomal proteins in Clusters 1, 2, 3 and Clusters A, B, C

